# Integrated multi-omics analysis to study the effects of simulated weightlessness on rhesus macaques *(Macaca mulatta)*

**DOI:** 10.1101/513382

**Authors:** Peng Zhang, Libin Shao, Jie Zhang, Wenjiong Li, Guangyi Fan, Ying Zhou, Guanghan Kan, Hongju Liu, Weidong Li, Fei Wang, Xixia Chu, Peng Han, Ling Peng, Xingmin Liu, Jianwei Chen, Xinming Liang, Jingkai Ji, Shiyi Du, Zhanlong Mei, Ronghui Li, Xun Xu, Shanguang Chen, Xin Liu, Xiaoping Chen

## Abstract

Safety and health of astronauts in space is one of the most important aspects of space exploration, however, the genomic research about how a weightless space can affect astronaut’s health was limited. In this study, we sequenced 25 transcriptomic, 42 metabolomic and 35 metagenomic data of 15 rhesus macaques *(Macaca mulatta)* spanning seven simulated weightlessness experiment stages. We identified 84 genes, 1911 features and 55 genera which are significantly changed in blood and muscle, hippocampal region, dorsomedial prefrontal cortex as well as fecal, respectively. Furthermore, performing the integrated analysis of three omics data, we found several pathways which were related to regulation of immune system process, glucose uptake, reaction to threatens, neurotoxic and bone or joints damage, such as tyrosine metabolism and tryptophan metabolism. Our results provided an initial attempt of “multi-omics” approaches which combined transcriptomics, metabolomics and metagenomics to illustrate some molecular clues for simulated weightlessness effect on the rhesus macaques and potential sight of microgravity’s effect on astronauts’ health.

## Introduction

Since the first traveling to space, the frequency of long-term spaceflight has increased rapidly. However, health issues are threatening mankind’s space exploring and the normal life of retired astronauts. Because research under real condition of spaceflight is limited, researchers often use ground-based analogs to study spaceflight’s effect on organisms. One such popular model is the head down-tilt bed rest (HDBR) which mimics the headward fluid shift and axial body unloading of spaceflight [1], and in which organisms remain either horizontal or −6 degree HDBR for days to months [1, 2]. Several previous studies suggested that long duration space flight can affect intracranial pressure [3], brain structure and function [4, 5], osteoclastogenesis [6], blood pressure [7], visual impairment [8] and the immune system [9, 10], indicating that solutions for maintain astronaut’s health. Multiple studies on space traveling astronauts, animals and plants, as well as microgravity analogues on animal models have been taken to study the biology changes during and after the spaceflight [11, 14].

The characteristics of space environment have two main factors, space radiation and microgravity [15]. It has been reported that the space radiation can damage DNA, leading to potentially harmful health consequences [15]. Space flight, especially microgravity, is one of the most extreme conditions that humans encounter [16]. However, little is known about the changes of gene expression, gut microbiota and metabolites of astronaut under microgravity. Addition of omics profiling to microgravity space flight experiments will understand key areas of variance in the molecular landscape [17]. The genomic and transcriptome studies can find gene expressions under difference environmental [18]. Space flight affects the community behavior of bacteria that harmful and beneficial human microbial interactions change during space flight [19]. About microgravity of systemic immune dysfunction may make the host more susceptible to pathogen infection [16]. Besides, metabolomics, the comprehensive study of metabolic reactions, was applied to study the metabolite profile in the studies of spaceflight effects on astronauts [2, 7], which help to understand the mechanism of physiological changes during and after space life. Thus, the results of integrated omics analysis could be used for aerospace medicine and research [17].

In this study, we recruited 15 rhesus macaques in different stages of ground-based HDBR analog using transcriptomics, metabolomics and metagenomics to understand the effects of weightless space flight analogues (HDBR) in rhesus macaques. As far as we know, our study will be the first multi-omics sights into the molecular study of space flight analogues.

## Results

### Metabolism significance of rhesus macaques in HDBR study

To investigate the effect of spaceflight on metabolism, we established a head down-tilt bed rest (HDBR) model using rhesus macaques. We then used ultra-performance liquid chromatography / mass spectrometry (UPLC/MS)-based metabolomics approach to quantify the metabolism signatures of the muscle, hippocampal region (HIP), dorsomedial prefrontal cortex (dmPFC) of 15 rhesus macaques in three different time points: pre-HDBR (control, T1), HDBR (T4) and recovery states (T7) (Figure 1, Table S1). We firstly conducted pair-wised comparison among these three time points using univariate and multivariable analysis. 356, 287 and 100 significantly different abundance features (DAFs) were identified in muscle by comparing HDBR to control, recovery to HDBR, and recovery to control, respectively (P < 0.05, fold change > 1.2 or fold change < 1/1.2, VIP>1, Figure S1). Besides, we also identified 328, 430 and 177 DAFs in HIP; 368, 472 and 109 DAFs in dmPFC. For the further analysis, we mainly focused on the 154, 198 and 207 DAFs that were significantly changed in HDBR comparing to control and recovery in muscle, HIP and dmPFC, respectively (P<0.05, Figure 2a, Figures S2a and S2b). Based on KEGG pathways analysis (Table S5), we found that the tryptophan metabolism pathway emerged in all comparisons of three tissues and tyrosine metabolism pathway contained 8 metabolites emerged in comparisons of muscle and dmPFC. Out of these eight metabolites, L-dopa (3,4-dihydroxy-L-phenylalanine, C00355, HMDB00609), L-Norepinephrine (C00547, HMDB00216), epinephrine (C00788, HMDB00068), L-Metanephrine (C05588, HMDB04063) were reported to be involved in neurotransmitter precursor, blood pressure control, stress reaction, glucose uptake and energy metabolism [20, 24].

**Figure 1.**
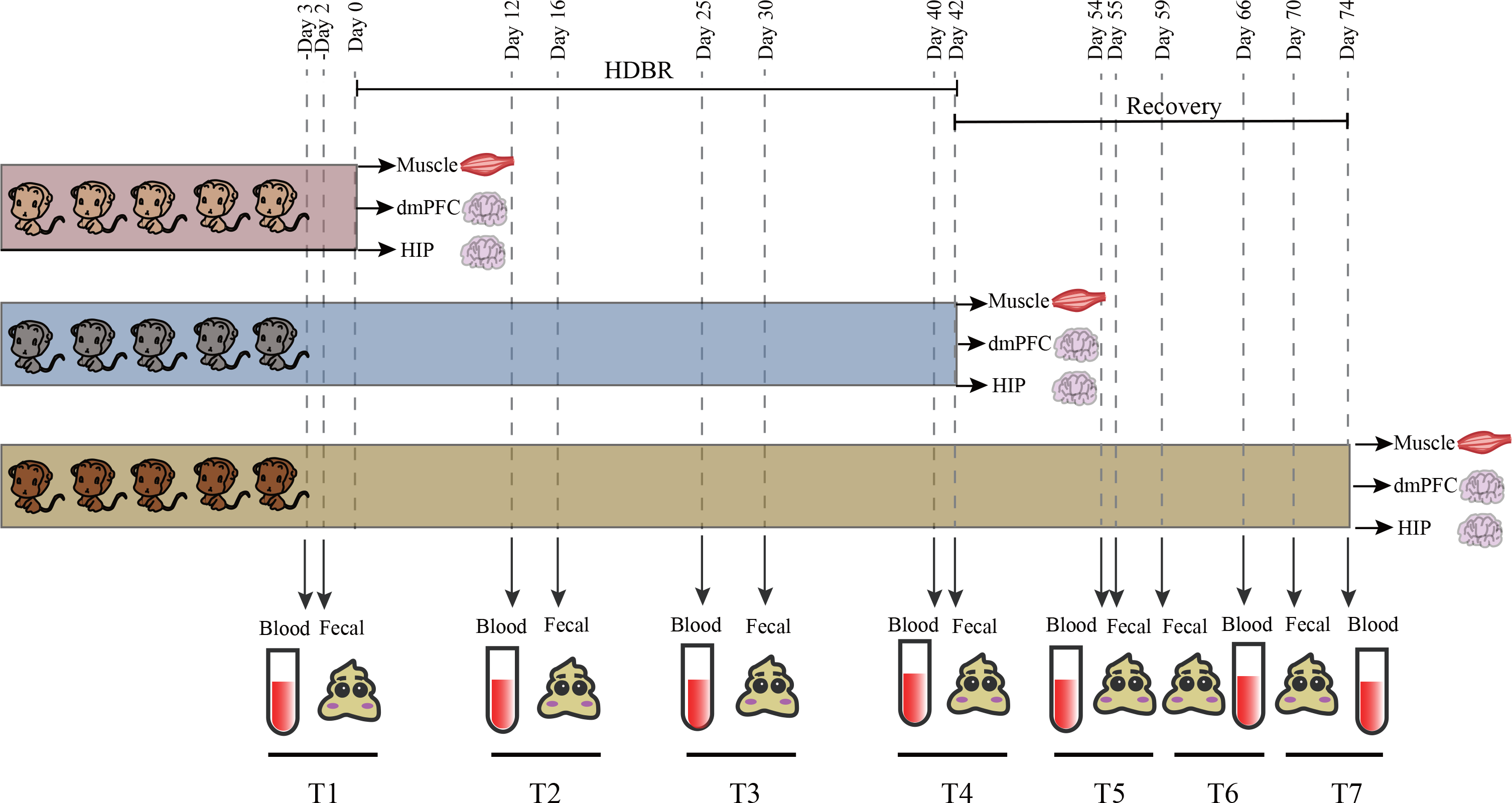
Overview of the study integrating metabolome, metagenome and transcriptome data and rhesus macaques HDBR experiments. Experimental Design for 15 rhesus macaques HDBR experiments. Each group of 5 rhesus macaques participated in this experiment. 5 rhesus macaques before HDBR were chosen for muscle, HIP and dmPFC tissues obtaining. 5 of the others took part in following HDBR experiment, then muscle, HIP and dmPFC tissues of which at HDBR 42 days were obtained. The 5 remaining rhesus macaques got into recovery experiments, and muscle, HIP and dmPFC tissues were obtained at 32 days after recovery experiment. Among all the experiments, blood samples 3 days before HDBR, 12 days, 25, and 40 days during HDBR, 12 days, 24, 32 days during recovery, and fecal samples 2 days before HDBR, 16 days, 30, and 42 days during HDBR, and 13 days, 17 and 28 days during recovery were all collected from the last 5 rhesus macaques.

**Figure 2.**
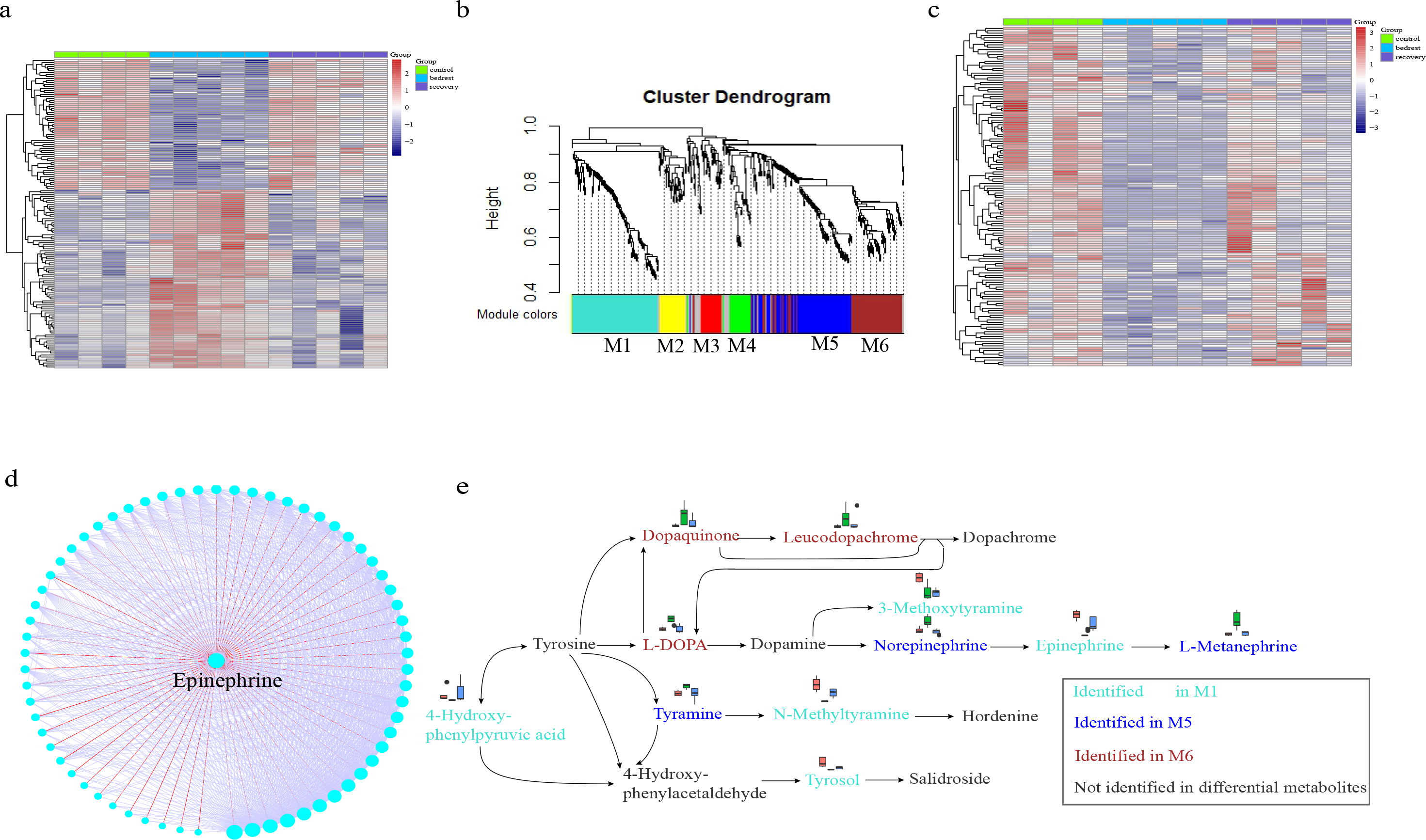
Meaningful metabolites and tyrosine metabolism pathway were found in muscle by metabolomics analysis. (a)A heatmap of 154 DAFs which only changing significantly in HDBR comparing to control and recovery in muscle.(b)Dendrogram of 552 muscle DAFs mainly clustering into 6 modules by WGCNA analysis. 6 modules colored with ‘turquoise’, ‘yellow’, ‘red’, ‘green’, ‘blue’ and ‘brown’ were labeled by ‘M1’ - ‘M6’, respectively. And ‘grey’ module included the remaining DAFs that did not fit clustering criteria.(c)A heatmap of DAFs in M1 showing that all DAFs down-regulated during HDBR and up-regulated during recovery in muscle.(d)Epinephrine was the hub metabolite of M1 network in muscle. Correlation of M1 DAFs were calculated by WGCNA. And the bigger size of the node means the more neighbor of this node. The adjacent lines of epinephrine colored with ‘red’, and the others colored with ‘light gray’.(e)11 significantly changed muscle metabolites showing in tyrosine metabolism pathway. 5 metabolites in M1 decreasing during HDBR and increasing during recovery were colored with ‘turquoise’, 3 metabolites in M2 showing opposite changes to M1 were colored with blue. And 5 metabolites in M3 showing similar trend with M1 were colored with brown. The 5 remaining metabolites colored with grey weren’t found in differential metabolites.

In addition, to observe the interaction of these DAFs, we performed the weighted correlation network analysis (WGCNA) for three tissues, respectively [34]. In details, we classified 552 DAFs into six modules in muscle, 668 DAFs into seven modules in HIP and 691 DAFs into six modules in dmPFC, respectively (Figure 2b, Figures S3a and S3b). Particularly, we found the DAFs of tyrosine metabolism pathway (mcc00350) mainly distributed in three modules (M1, M5 and M6). Interestingly, the DAFs of M1 showed a good co-abundant pattern in muscle that all of these features decreased in HDBR and increased in recovery (Figure S3c and Figure 2c). Among them, the abundance of L-Alanine (C00041, HMDB00161), L-Carnitine (C00318, HMDB00062) and calcitroic acid (HMDB06472) were decreased, which might be related to the muscle weakness and bone loss (Figures S3d-S3f). L-Alanine (C00041, HMDB00161) is an important amino acid in glucose-alanine cycle, which moves gluconeogenesis from muscle to liver to produce ATP stored in muscle for muscle contraction [26]. L-Carnitine ((C00041, HMDB00161) is most abundant in skeletal muscle and cardiac muscle, and it actives and transports fatty acids into the mitochondria for energy generation in skeletal muscle [27]. Calcitroic acid (HMDB06472) is a major metabolite from 1,25-Dihydroxyvitamin D3 which plays an important role on bone homeostasis and resorption, was found decreasing after long-term bed rest or space flight [38] [28]. Besides, we found that epinephrine (C00788, HMDB00068) was increased the metabolic rate in skeletal muscle (C00788, HMDB00068) [30], and it was the hub metabolite of M1 (Figure 2d). Pyridoxamine (C00534, HMDB01431) was the hub metabolite of M6, which was one form of vitamin B_6_ taking part in metabolism of L-Alanine.

### Gut microbiome changes of rhesus macaque in HDBR study

The gastrointestinal tract harbors most of microbes, which play important roles in organisms’ health, disease, immunity, and even behavior [31, 32]. To investigate the effect of HDBR on the microbiome, we sequenced 35 metagenomics gut samples from five rhesus macaque individuals of seven time points (T1~T7) of HDBR analog (Figure 1). We generated a total of 286.86 Gb raw data and 275.99 Gb high-quality data after removing low quality data and the host, the average clean data was 7.89 Gb for each sample (Table S2). Then, we constructed a reference gene catalogue of rhesus macaque’s gut metagenome using all samples (Table S3). Among of these genes, 63.10% of gene catalogue could be annotated to phylum level, 21.00% to genus level, and 2.43% to species level. And 45.51% could be annotated to 6,631 KEGG Orthologs (KOs).

From the annotation of microbiome species, we found that Firmicutes and Bacteroidetes were dominated on the phylum level, as well as Prevotella and Clostridium were the dominated genera on the genus level (Figure S4). From the annotation of KEGG function, K03088, K03327, K06147, K00599 and K00754 were most dominated KOs with a higher abundance which annotated to transcription machinery, ion-coupled transporters, transporters, histidine metabolism/tyrosine metabolism and fructose and mannose metabolism. In all, 49.48% of genera and 95.58% of KO functions were shared by monkey, human gut gene catalogues (Figure S5).

We identified 55 genera with a significant change of abundance from T1 to T7 (Table S6). Interestingly, out of these 55 genera, 43% of them had decreased in their abundance in the experimental stages (T2~T4) and increase during the recovery phases (T5-T7). These genera include such as *Acinetobacter* and *Lactococcus*, which is involved regulation of inflammation [33, 34, 35] and protection of infections [36], indicating that HDBR caused a disorder in the rhesus macaque’s intestinal flora (Figure 3a and 3b). The abundance of two genera had been decrease continuously throughout the study, such as *Bifidobacterium*, which could modulate host immune responses, inhibit infection by pathogens, and regulate intestinal microbial homeostasis [37, 38, 39] (Figure 3c). In addition, we found 537 KOs with significantly changed abundance were assigned into 139 genera, of which 27 genera were significantly changed during T1 to T7 (P<0.05). Correlation analysis showed that 44.76% of pair-wise correlation between KOs and genera were positive, whereas only 3.38% correlation were negative (R2>0.3, P<0.05) (Figure 3d). This results indicate that the changes of genera abundance positively affect the gut microbial function, such as *Myroides* and *Acinetobacter*, which could help to improve the K00121, K00151, K00276, K00451 and K01555 which participate in tyrosine metabolism pathway. In summary, our findings revealed that HDBR affects the abundance of gut microbiome, which might be related to the host immune response and metabolism.

**Figure 3.**
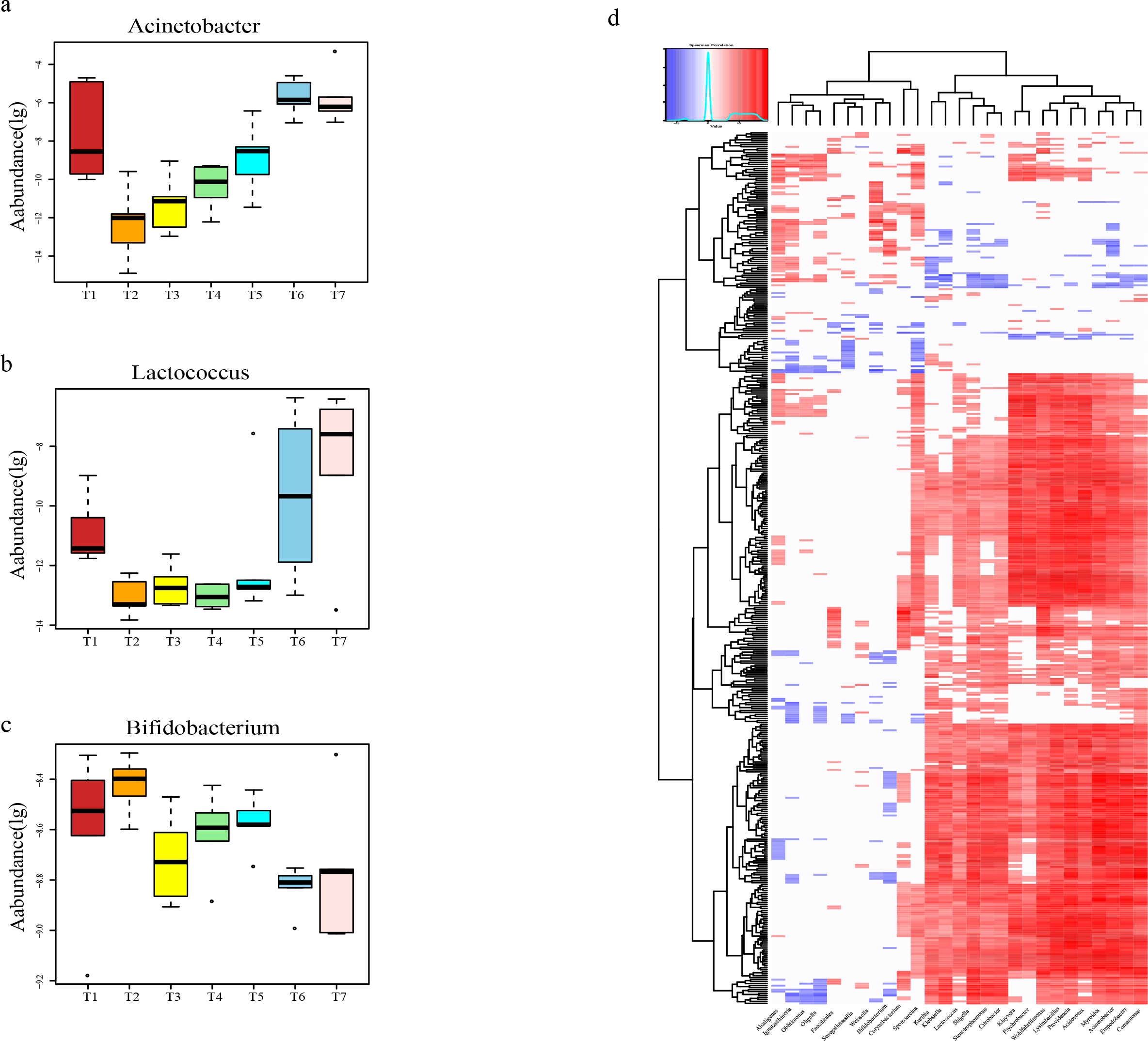
Several important intestinal genera alterations in seven time points with 95% confidence interval for the medians, and associations of differential gut microbial genera with differential KOs. (a-c) Box plot of the differential genera abundance of Acinetobacter, Lactococcus and Bifidobacterium. The x coordinate represents T1～T7 time points, y coordinate represents the genera abundance (log10). For each interquartile ranges (IQRs), the first and third quartiles were showed as the boxes, and the line inside the box represents the median. The circles data points represent the outside of the whiskers which marked as the lowest or highest values within 1.5 times IQR boxes.(d) Spearman correlation coefficient heatmap (P<0.05) between 27 genera differed in abundance (x coordinate) and 442 Kos differed in abundance (y coordinate). The red color represented significantly positive correlation, the blue color represented significantly negative correlation, and blank areas indicate no significant correlation.

### Transcription features of rhesus macaque in HDBR study

To investigate the effect of HDBR on gene expression in immune cells, we collected 25 blood samples from five rhesus macaque individuals of six time points (T1, T3~T7) (Figure 1). We generated 779.09 M reads in six time points of HDBR analog (Table S4). Using pairwise comparison of each time point, we detected 84 significantly differential expressed genes (DEGs, fold change>2 and P<0.05) in at least two time points. We firstly focused on 65 DEGs in HDBR (T3 and T4) comparing to the control (T1) and the recovery phases (T5~T7, Figure 4a). These DEGs were significantly enriched in 44 biological processes (P<0.05), and 41 of them were related to the regulation of immune system, including regulation of leukocyte activation (GO:0002694), regulation of T cell differentiation (GO:0045580) and regulation of interleukin-2 production (GO:0032663) (Figure 4b and Table S7). DEGs in these immune response biological processes were mainly down-regulated in the HDBR, which were consistent with the previous researches about the immune system of animal and human in simulated long-term microgravity environment [36][40]. In addition, we also found 71 DEGs in recovery status (T5~T7) comparing to the control (T1, Figure 4a), which were also mainly enriched in the immune system. Besides, WGCNA [25] method was employed to construct genes co-expression networks in transcriptomes, 1941 genes with max median absolute deviation (MAD) were clustered into eleven modules. In accordance with the differential expression analysis, genes in the purple module were mainly related to immune response (Table S8) with the expression dropped in HDBR.

**Figure 4.**
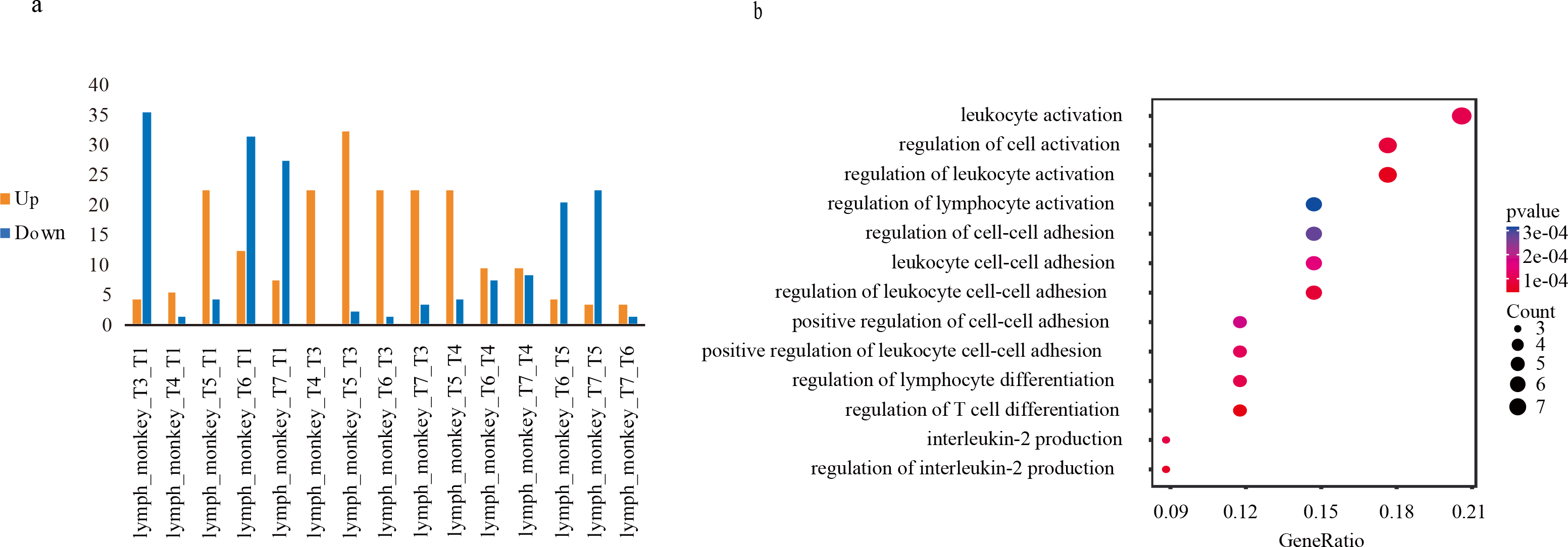
Interesting DEGs and biological processes found in transcriptomics analysis. (a)Histograms of DEGs between any two time points. Up-regulated and down-regulated DEGs were colored with ‘orange’ and ‘blue’, respectively. (b)GO analyses of the 65 DEGs showing a significant enrichment of several biological processes.

### Integrated analysis of omics data in rhesus macaque

Short-chain fatty acids (SCFA) is a dominant metabolite produced from bacteria, related to the immune response, such as butyrate regulates the size and function of the regulatory T cell network by promoting the induction and fitness of regulatory T cells in the colonic environment [41, 42, 43]. In our study, DEGs in lymphatic cells were related to reduce immune response and dysregulation of muscle butyrate metabolism. In the gut microbiome, the relative abundance of butyrate-producing colon bacteria *Eubacterium*, *Roseburia* and their cross-feeding bacteria *Bifidobacteria* were reduced in the HDBR. Thus, our results revealed that the weak immune response and the reduced abundance of butyrate during HDBR might be related to the abundance change of gut microbe (Figure 5).

**Figure 5.**
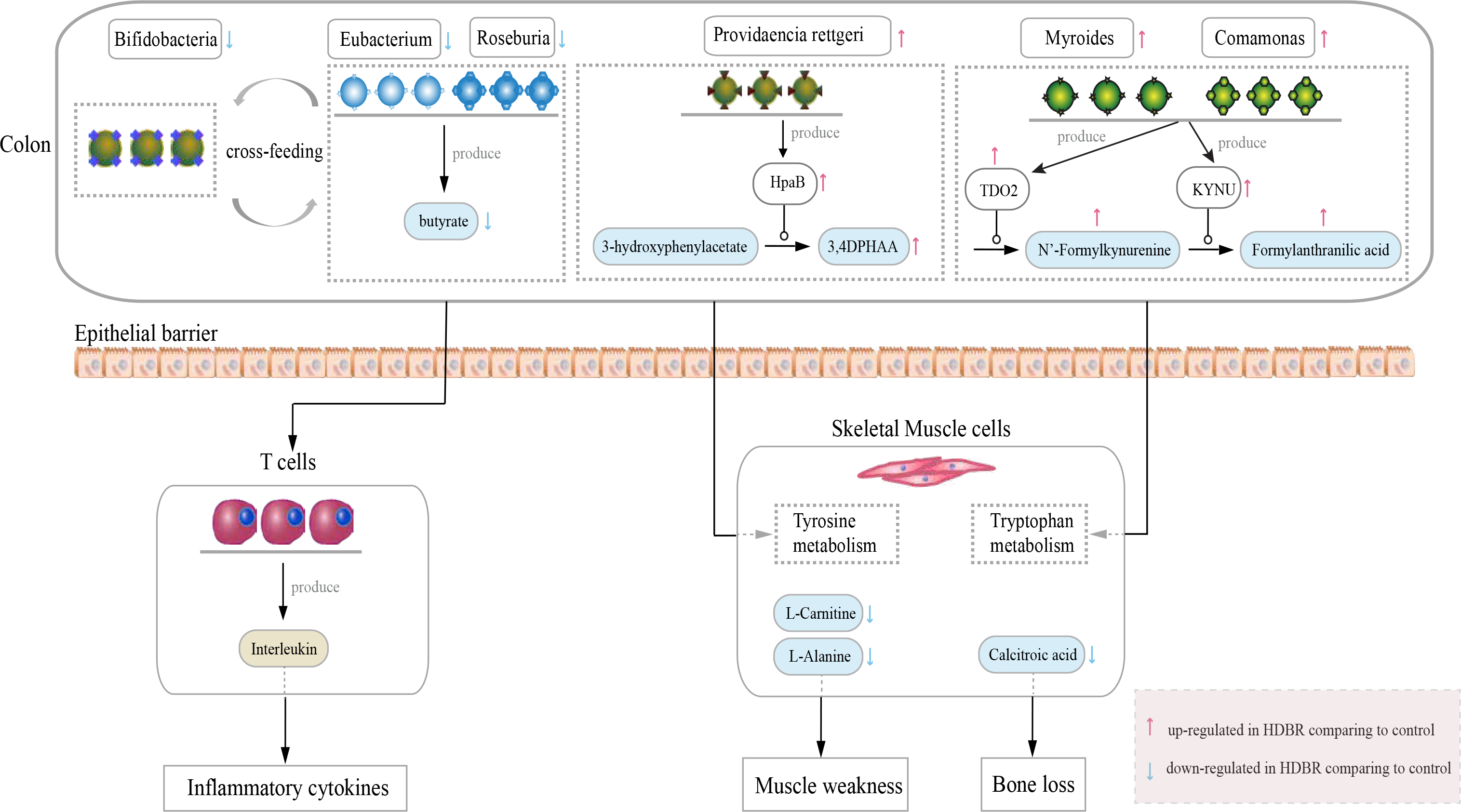
Reducing gut microbes produced lesser butyrate to weaken the host immune response, and the gut microbes might take part in tyrosine metabolism (mcc00350) and tryptophan metabolism pathway (mcc00380).

Besides, previous study showed that 3-hydroxyphenylacetate is a by-product of the tyrosine metabolism mediated by *Clostridium spp* [28]. Enzyme 4-hydroxyphenylacetate 3-monooxygenase (EC: 1.14.14.9), which was not found in rhesus macaque, could transform 3-hydroxyphenylacetate (C00642, HMDB00020) to 3,4-Dihydroxybenzeneacetic acid (3,4DPHAA, C01161, HMDB01336) in tyrosine metabolism. Coincidently, enzyme 4-hydroxyphenylacetate 3-monooxygenase and 3,4DPHAA both increased in during HDBR and decreased during the recovery phase, consistent with the abundance changes of a gut microbe *Providencia rettgeri* which produces enzyme 4-hydroxyphenylacetate 3-monooxygenase (EC: 1.14.14.9), indicating *Providencia rettgeri* influence tyrosine metabolism by producing 3,4DPHAA (Figure 5). Besides, we also found enzyme tryptophan 2,3-dioxygenase (EC: 1.13.11.11) could produce N’-Formylkynurenine (C02700, HMDB60485), which then transformed to Formylanthranilic acid (C05653, HMDB04089) by enzyme kynureninase (EC: 3.7.1.3) in tryptophan metabolism. Enzyme tryptophan 2,3-dioxygenase and kynureninase could be produced by the gut microbes with abundance change during HDBR, such as *Myroides* and *Comamonas*. Furthermore, we found *Myroide* were positively and significantly correlated with Leucodopachrome (C05604, HMDB04067) and Dopaquinone (C00822, HMDB01229) in the tyrosine metabolism pathway (mcc00350) and *Providencia* were found positively and significantly correlated with p-Cresol (C01468, HMDB01858) which is a metabolite of tyrosine (P<0.05). Meanwhile, *Lactococcus* were positively correlated with L-Carnitine (C00318, HMDB00062) which is crucial in providing energy to muscles [27] (Figure S6). Therefore, the significantly changed abundance of gut microbes were involved in tyrosine metabolism (mcc00350) and tryptophan metabolism (mcc00380) pathways during the HDBR analog (P<0.05).

## Discussion

Living in a space environment with microgravity and motionless for long periods of time may have adverse effects on immunity, metabolism and health. The first report from the Soviet immunologist, Konstantinova and coworkers [44], found that lymphocyte responsiveness to mitogens was remarkably reduced after astronauts after a longtime spaceflight. In this study, we applied transcriptomic, metabolomic and metagenomic analysis in a head down-tilt bed rest model to elucidate the effect of spaceflight and analogue microgravity environment on biology functions. In transcriptome data, DEGs were enriched in multiple biological processes mainly related to immune response, indicating that gene expression has plausible functional connections with spaceflight and microgravity. For the first time, we also found the effects of an analogue microgravity environment on immunity by combining metabolomics and metagenomics. We found that tyrosine metabolism, which plays an important role in muscle function, was affected by HDBR. The abundance of L-Dopa, norepinephrine, epinephrine and L-Metanephrine in catecholamines biosynthesis and metabolism pathway, dopaquinone and leucodopachrome in L-Dopachrome biosynthesis pathway and 4-Hydroxyphenylpyruvate and homogentisate in homogentisate biosynthesis pathway were significantly affected by HDBR (Figure 2e). We also found that calcitroic acid (related to bone balance), L-Alanine and L-Carnitine (related to muscle contraction and muscle weakness) were affected by HDBR.

To our knowledge, this was the first study successfully combining metabolomics and metagenomics in studying simulated microgravity on health and metabolism. The findings provided valuable insights into how long-term spaceflight could affect metabolism and health. More importantly, our findings on the link between gut microbiome with metabolism and immunity provided the possible solutions to combat dysregulated immunity and metabolism in spaceflight. This study was conducted with animal model in a simulated environment, we supposed a solid combination of omics in the study of astronauts could be a useful tool for future study of astronauts in extremely environment. Further study with more astronauts would provide more information during long term space flight.

## Materials and Method

### Animal experiments

We sampled 20 healthy male rhesus macaques, aged 4 to 6 years and weighing 4 to 8 kg from Beijing Institute of Xie’erxin Biology Resource (Beijing, China). All of these rhesus macaques received 3 months of domestication (involving preliminary caretaker handling, confinement jacket fitting, and tilt-table acclimation training) at the Laboratory Animal Center of China Astronaut Research and Training Center prior to the start of the experiments. Only 15 well-domesticated rhesus macaques were selected and separated into 3 groups: 1) CON: ground-based controls, 2) HDBR: 42 days of HDBR, 3) REC: 42 days of HDBR plus 32 days of recovery.

A six-week head-down tilted bed rest (HDBR) experiment was performed on rhesus macaques to simulate weightlessness as described previously [45]. Briefly, rhesus macaques laid on beds, which were tilted backward 10 °C from the horizontal. The head-down monkeys wore the confinement jacket, which enabled them to be fixed to the bed. These rhesus macaques were housed one per bed in rooms with air temperature maintained at 23 ± 2°C and a standard 12:12 h dark–light cycle (lights were turned on at 8:00 a.m. and off at 8:00 p.m.). After six weeks of HDBR, each rhesus macaque was solely removed into stainless steel mesh cages to recovery for 32 days. Throughout the duration of the experiment, rhesus macaques received an intensive humanistic care. For instance, the rhesus macaques always had free access to food and water. Toys (such as the drum-shaped rattle, a Chinese traditional toy) were available all the time except during experimental procedures. The caretaker accompanied the rhesus macaques during the day time to help relieve anxiety. The general health condition of the rhesus macaques was also carefully monitored. All procedures were performed in accordance with the principles of the Association for Assessment and Accreditation of Laboratory Animal Care International (AAALAC), approved by Institutional Animal Care and Use Committee of China Astronaut Research and Training Center (ACC-IACUC-2014-001).

### Sample collection

All samples provided consent forms which were approved by BGI genomics Committee of Ethics. Under light ketamine sedation, sterile heparinized peripheral blood samples were obtained from femoral vein of the five rhesus macaques in REC group before (Pre-3, T1), during (HB-12, HB-25, HB-40, T2~T4) and after (R-12, R-24, R-32, T5~T7) the HDBR at 10:00 a.m. Peripheral blood mononuclear cells (PBMCs) were then collected by Ficoll-Hypaque density-gradient centrifugation. Fecal samples were also collected from the five rhesus macaques in REC group before (Pre-2, T1), during (HB-16, HB-30, HB-42, T2~T4) and after (R-13, R-17, R-28, T5~T7) the HDBR. Skeletal muscle sample, dmPFC sample and HIP sample were collected from all of the selected rhesus macaques in each group.

### Transcriptome analysis

mRNA of the PBMCs was isolated by oligo(dT) and sequenced by Complete Genomics(CG) SE50, we then use SOAPnuke to filter out the reads of low quality, at least 20 million reads were generated for each sample. HISAT2 was applied to map sequence reads to genome and RSEM to calculate the FPKM. Spearman correlation between all samples were calculated using FPKM, the correlation between all samples was very high (>0.95) and the biological replicates did not cluster well together. We used DESeq2 package in R to find different expressed genes [46], and GO was used to annotated function information.

### Metabolomics analysis

UPLC-MS technology was implemented for metabolites detection. The experimental quality was evaluated by quality control (QC) samples. Features alignment, picking and identification were performed by Progenesis QI (Waters, Nonlinear Dynamics, Newcastle, UK). MetaX software was used for data cleaning and statistical analysis [47]. Low quality features were removed within data cleaning. By combining the univariate and multivariate statistical analysis, significantly changed features (P value<0.05, fold change <1/1.2 or fold change >1.2, VIP >1) were acquired. Those features were further annotated by Progenesis QI with Human Metabolome databases (version 3.6), and online Kyoto Encyclopedia of Genes and Genomes database (www.genome.jp/kegg/).

Clusters of co-abundant metabolites of muscle, HIP and dmPFC were performed by R package WGCNA [25], respectively. Soft threshold β=9 for muscle features, soft threshold β=13 for dmPFC features, and soft threshold β=12 for HIP features were chosen by scale free topology analysis, for signed, weighted features co-abundance correlation network construction. Dynamic hybrid tree-cutting algorithm by deepSplit of 4 and a minimum cluster size of 30 were applied for clusters identification. If the biweight mid-correlation between the cluster’s eigenvectors exceeded 0.8, the similar clusters would be subsequently merged. The muscle, dmPFC and HIP features clusters, were labelled by M1-M6, D1-D7 and H1-H6, respectively.

### Metagenomics analysis

Raw reads sequenced on the Illumina Hiseq 2000 platform (Expression Analysis Inc., San Diego, CA, USA) at BGI were filtered to remove the adaptor contamination, low-quality reads and host genomic DNA (Rhesus macaques, assembly Mmul_8.0.1, NCBI). The remaining high-quality reads were assembled by metaSPAdes (v3.10.1). Open Reading Frames (ORFs) in contigs of each sample were obtained using GeneMark (v2.7). The non-redundant gene set of all ORFs was clustered using CD-HIT (v4.5.7) based on nucleotide sequence and the identity is 95% at the coverage 90%. Taxonomic annotation of gene set was made with CARMA3 on the basis of BLASTP alignment with bacteria and archaea from NCBI-NR database. And the gene set was also annotated against KEGG (v59) databases with BLAST (v2.2.23).

Gene abundance profiling was calculated based on the alignment of SOAP2 and the species abundant profile and functional profile were summarized from their respective genes [48]. The differential alpha diversity, species and KOs of different time points were calculated using R kruskal.test. Spearman coefficient was used to calculate the relationship between different genus and different Kos, between different genus and different metabolic features according to their abundance.

## Supporting information

none

none

## Acknowledgments

This work was supported by grants from the National Basic Research Program of China (2011CB711003), the National Natural Science Foundation of China (81772016, 81272177) and the 1226 major project (AWS16J018) to X.C.

