## Supplementary material for "Integrated multi-omics analysis to study the effects of simulated weightlessness on rhesus macaques *(Macaca mulatta)*": none

**Supplemental Information**

**Integrated multi-omics analysis to study long term microgravity**

**effect on *Macaca mulatta***

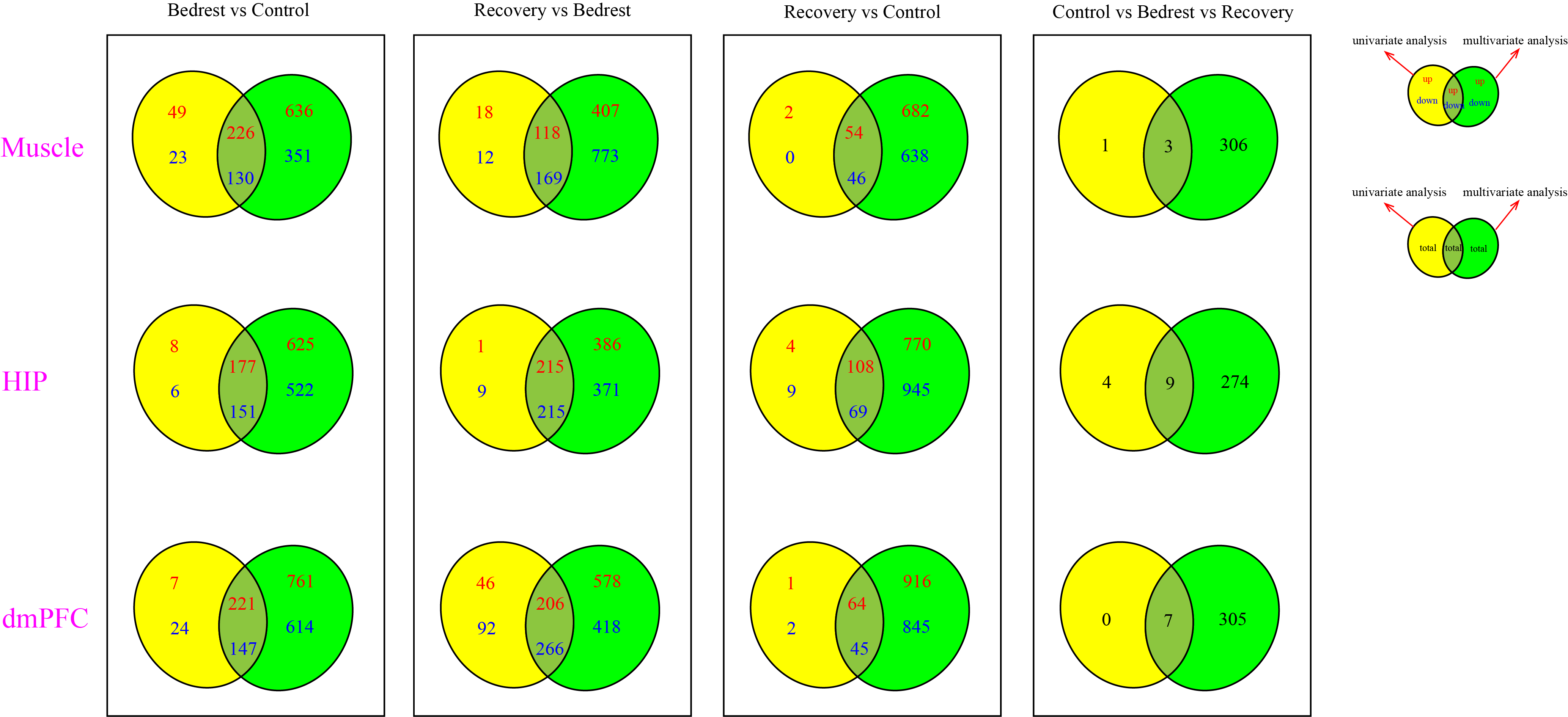

**Figure S1, related to figure 2. Venn diagrams of different feature detected by univariate and multivariate analysis in muscle, HIP and dmPFC, respectively.**

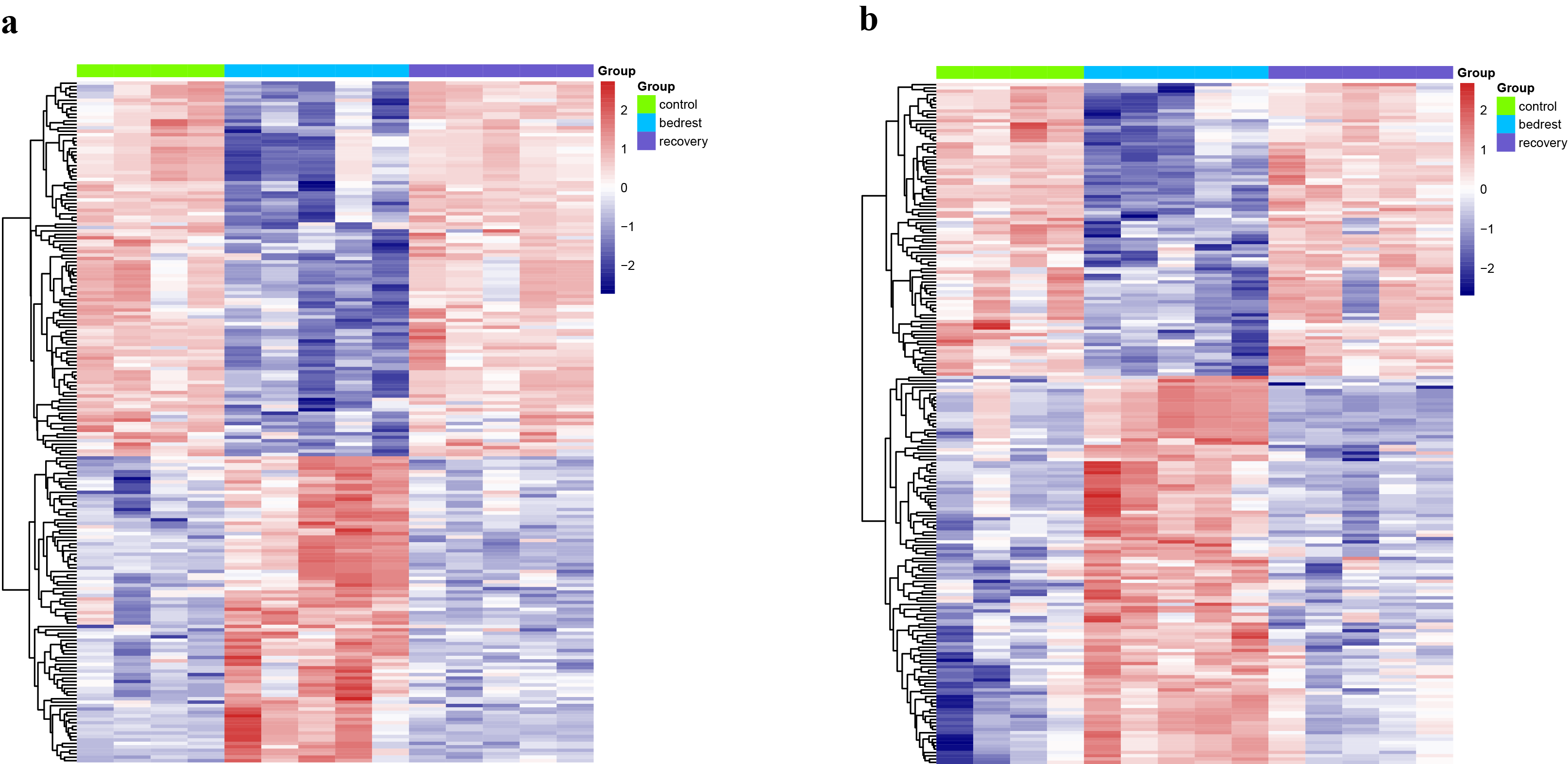

**Figure S2, related to figure 2. Heatmaps of DAFs which only changed significantly in HDBR comparing to control and recovery in HIP and dmPFC samples, respectively.**

(a)	A heatmap of DAFs that only changed significantly in HDBR comparing to control and recovery in HIP samples.

(b)	A heatmap of DAFs that only changed significantly in HDBR comparing to control and recovery in dmPFC samples.

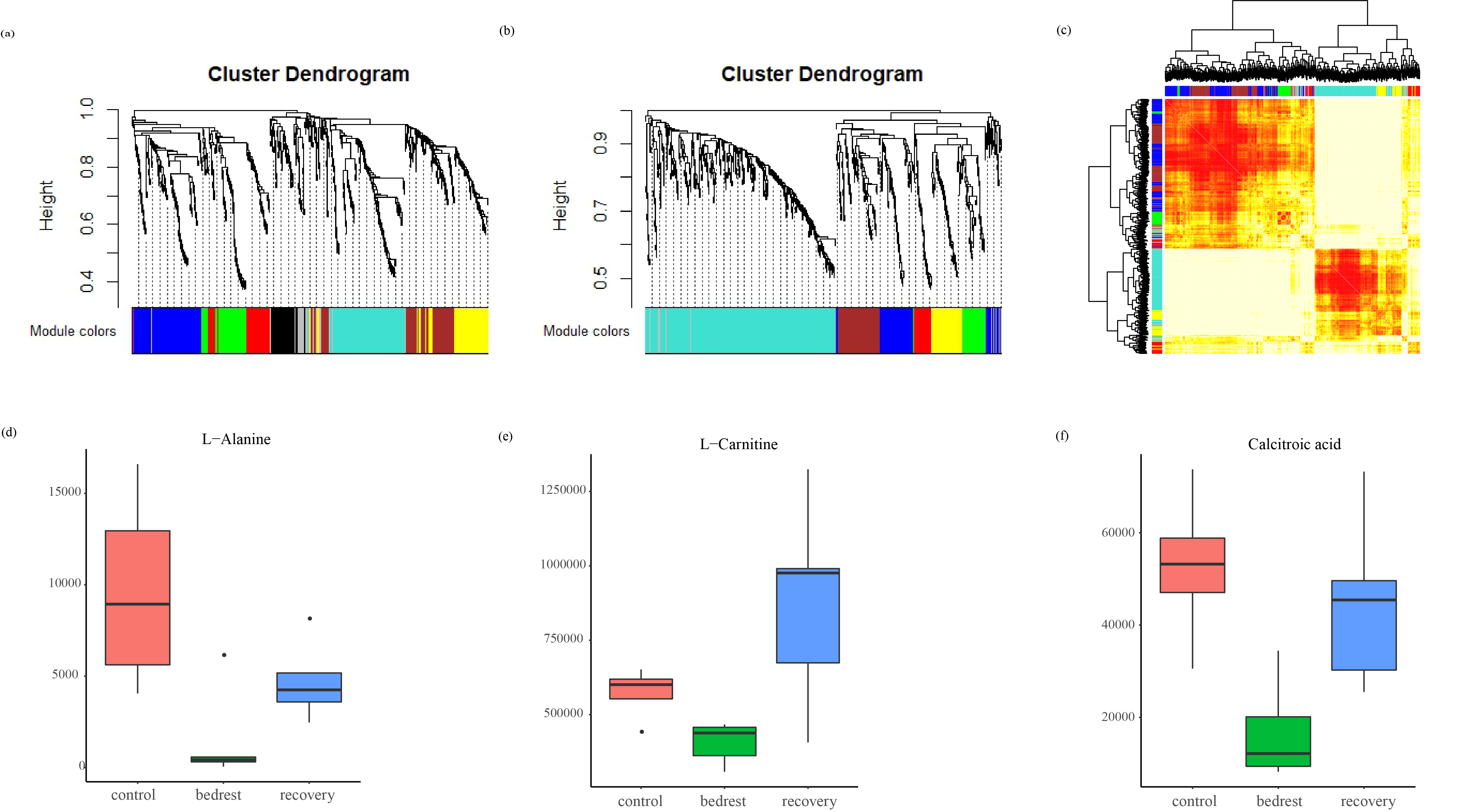

**Figure S3, related to figure 2. Modules in HIP and dmPFC, M1 showing a good co-abundance in muscle, and the abundance of three interesting muscle metabolites in three time points.**

(a)	Dendrogram of 668 HIP DAFs mainly clustering into 7 modules by WGCNA analysis. ‘Grey’ module included the remaining DAFs that did not fit clustering criteria.

(b)	Dendrogram of 691 dmPFC DAFs mainly clustering into 6 modules by WGCNA analysis. ‘Grey’ module included the remaining DAFs that did not fit clustering criteria.

(c)	Correlation of all muscle DAFs showing a good similarity in M1. The DAFs corresponded to rows and columns. 6 modules colored with ‘turquoise’, ‘yellow’, ‘red’, ‘green’, ’blue’ and ‘brown’ represent ‘M1’ - ‘M6’, respectively. And ‘grey’ module was the remaining DAFs that did not fit clustering criteria.

(d)	Abundance of L-Alanine in muscle during control, bedrest and recovery, respectively. L-Alanine decreased during bedrest, then increased during recovery.

(e)	Abundance of L-Carnitine in muscle during control, bedrest and recovery, respectively. L-Alanine decreased during bedrest, then increased during recovery.

(f)	Abundance of calcitroic acid in muscle during control, bedrest and recovery, respectively. L-Alanine decreased during bedrest, then increased during recovery.

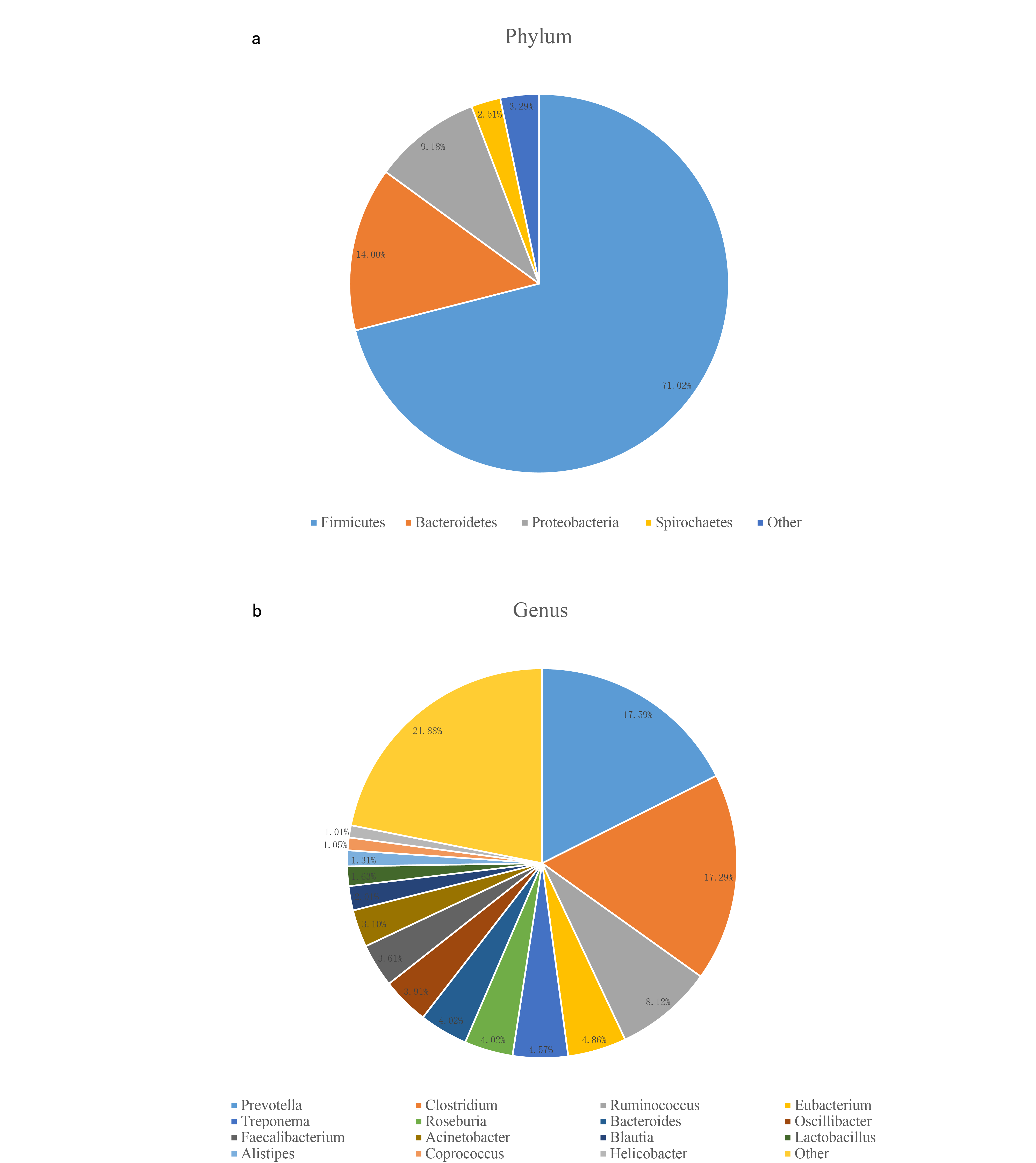

**Figure S4, related to figure 3. Taxonomic annotation of the 3.2M monkey gut gene catalogue.**

(a)	Denoted the proportion of the 5 most abundant phyla.

(b)	Denoted the proportion of the 16 most abundant genera.

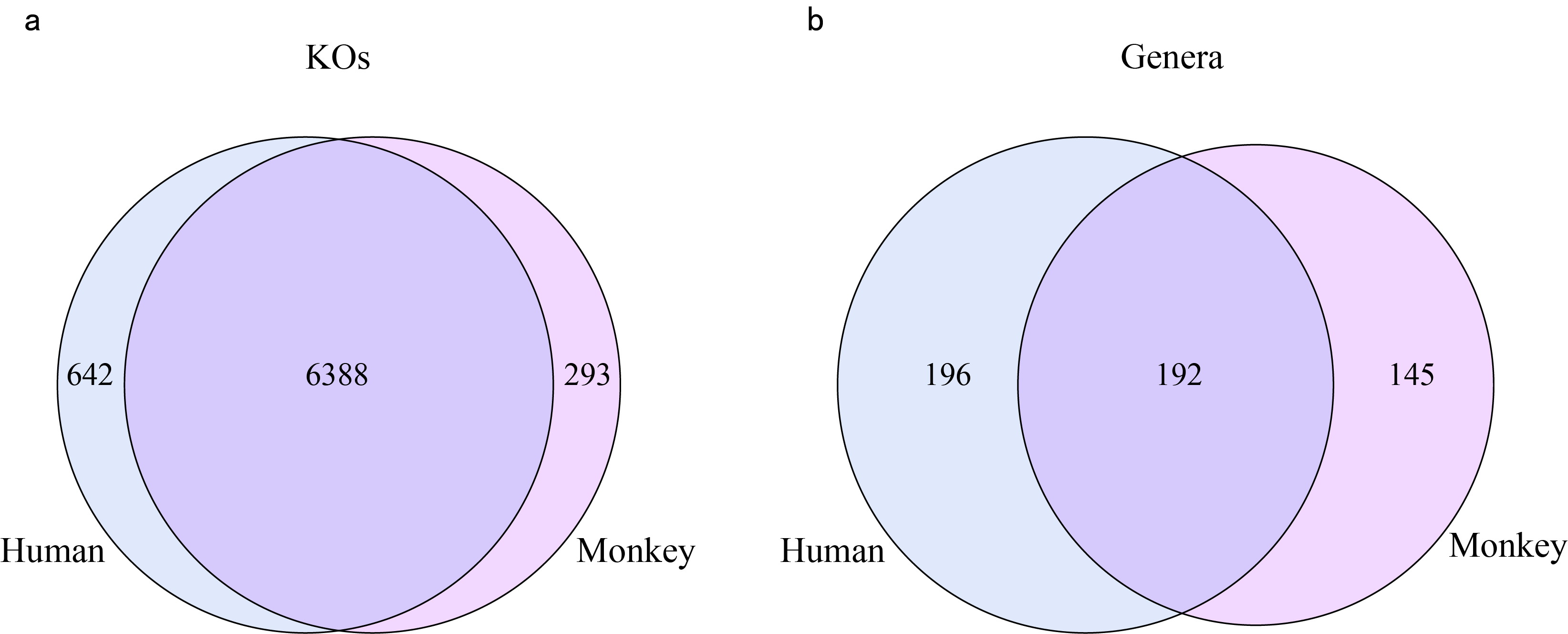

**Figure S5, related to figure 3. Venn diagram of gut microbial KO functions and genera between monkey and human.**

(a-b) Shared KO functions and genera across human and monkey gut microbial annotation. Each circle is the number of genes or the number of functions annotated by the gut microbe. The left represents the specifically of human gut, and the right represents the specifically of monkey gut. The overlap parts appear in both two species.

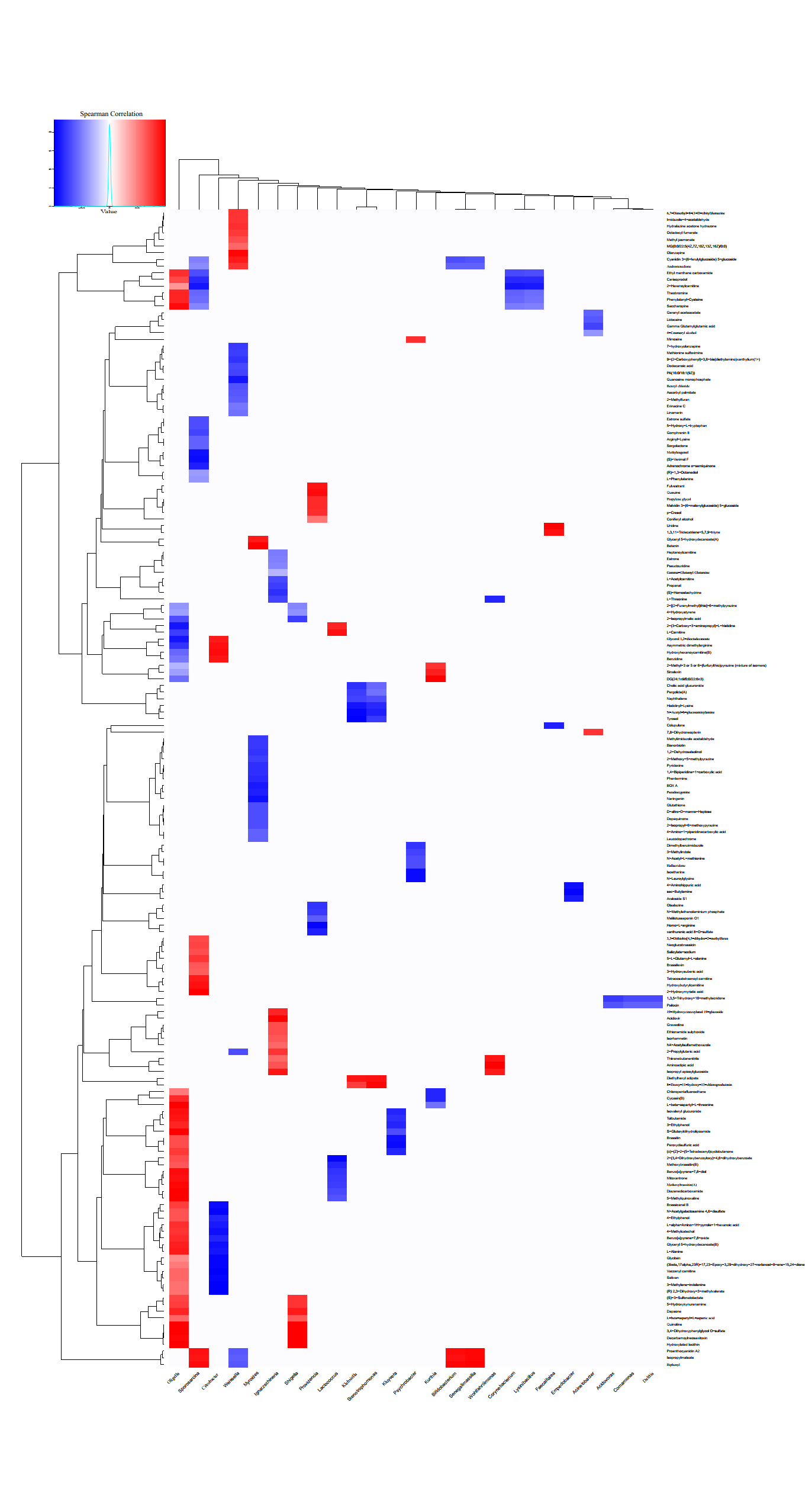

**Figure S6, related to figure 5. Associations of differential intestinal genera with differential muscle metabolites.**

Spearman’s rank correlation of T7 between 27 differential genera and differential muscle metabolites. The x coordinate represents 27 differential genera, y coordinate represents the 175 differential muscle metabolites with significant correlation. The red color represented significantly positive correlation and the blue color represented significantly negative correlation, the white color means no significant correlation.

|  | **Mode** | **Muscle** | **HIP** | **dmPFC** |
| --- | --- | --- | --- | --- |
| Raw Features | Positive | 8,105 | 8,109 | 8,109 |
|  | Negative | 3,950 | 5,339 | 5,339 |
| Clean Features | Positive | 5,550 | 6,581 | 6,581 |
|  | Negative | 2525 | 2,958 | 2,958 |

**Table S1, related to figure 2. Features number before and after quality control in muscle, HIP and dmPFC metabolomics, respectively.**

| **SampleID** | **Raw(Gb)** | **After_QC(Gb)** | **After_rmHost(Gb)** |
| --- | --- | --- | --- |
| M081533-140724A | 7.74 | 7.52 | 7.51 |
| M081533-140811A | 8.50 | 8.26 | 8.25 |
| M081533-140825A | 9.39 | 9.11 | 9.09 |
| M081533-140906A | 9.09 | 8.73 | 8.71 |
| M081533-140909A | 7.78 | 7.60 | 7.44 |
| M081533-140923A | 7.76 | 7.54 | 7.51 |
| M081533-141004A | 9.32 | 8.98 | 8.56 |
| M090341-140724A | 7.34 | 7.17 | 7.07 |
| M090341-140811A | 8.68 | 8.42 | 8.39 |
| M090341-140825A | 7.87 | 7.61 | 7.59 |
| M090341-140906A | 8.43 | 8.17 | 8.16 |
| M090341-140909A | 7.74 | 7.49 | 7.47 |
| M090341-140923A | 6.63 | 6.44 | 6.43 |
| M090341-141004A | 7.82 | 7.58 | 7.55 |
| M090997-140724A | 8.36 | 8.12 | 8.10 |
| M090997-140811A | 8.72 | 8.47 | 8.45 |
| M090997-140825A | 8.94 | 8.68 | 8.66 |
| M090997-140906A | 7.25 | 7.04 | 7.03 |
| M090997-140909A | 8.30 | 8.03 | 8.01 |
| M090997-140923A | 7.13 | 6.82 | 6.81 |
| M090997-141004A | 7.24 | 7.02 | 7.00 |
| M091289-140724A | 7.54 | 7.32 | 7.31 |
| M091289-140811A | 8.33 | 8.08 | 8.05 |
| M091289-140825A | 7.65 | 7.36 | 7.35 |
| M091289-140906A | 8.69 | 8.42 | 8.41 |
| M091289-140909A | 8.29 | 8.04 | 8.02 |
| M091289-140923A | 8.67 | 8.45 | 8.43 |
| M091289-141004A | 8.36 | 8.12 | 8.10 |
| M101043-140724A | 8.75 | 8.50 | 7.99 |
| M101043-140811A | 8.07 | 7.82 | 7.81 |
| M101043-140825A | 8.96 | 8.68 | 8.54 |
| M101043-140906A | 8.88 | 8.61 | 8.59 |
| M101043-140909A | 7.89 | 7.64 | 7.62 |
| M101043-140923A | 9.67 | 9.30 | 9.28 |
| M101043-141004A | 7.05 | 6.82 | 6.71 |
| Average | 8.20 | 7.94 | 7.89 |

**Table S2, related to figure 3. Sequencing data of metagenome before and after quality control, and after removing host data.**

| **SampleID** | **Number** | | **Length(Mp)** | | **N50(bp)** | | **N90(bp)** | | **GC(%)** | **MapRate(%)** | |
| --- | --- | --- | --- | --- | --- | --- | --- | --- | --- | --- | --- |
| M081533-140724A | 524,074 | 371.7 | | 861 | | 387 | | 48.51 | | | 74 |
| M081533-140811A | 414,298 | 297.05 | | 882 | | 387 | | 50.24 | | | 75.73 |
| M081533-140825A | 417,198 | 303.73 | | 900 | | 390 | | 49.7 | | | 77.91 |
| M081533-140906A | 441,689 | 322.36 | | 897 | | 396 | | 48.29 | | | 74.2 |
| M081533-140909A | 388,122 | 283.16 | | 894 | | 399 | | 48.79 | | | 74.63 |
| M081533-140923A | 345,962 | 261.16 | | 942 | | 405 | | 50.1 | | | 77.41 |
| M081533-141004A | 377,438 | 282.98 | | 939 | | 402 | | 50.33 | | | 78.39 |
| M090341-140724A | 437,230 | 302.88 | | 837 | | 375 | | 46.96 | | | 69.86 |
| M090341-140811A | 508,907 | 369.24 | | 894 | | 390 | | 50.24 | | | 75.9 |
| M090341-140825A | 494,227 | 357.49 | | 882 | | 393 | | 50.05 | | | 76.21 |
| M090341-140906A | 533,163 | 382.74 | | 879 | | 390 | | 49.46 | | | 76.12 |
| M090341-140909A | 506,916 | 372.74 | | 897 | | 405 | | 50.31 | | | 76.95 |
| M090341-140923A | 389,913 | 278.81 | | 867 | | 390 | | 48.03 | | | 77.25 |
| M090341-141004A | 455,319 | 333.19 | | 894 | | 396 | | 48.03 | | | 77.13 |
| M090997-140724A | 537,681 | 378.38 | | 855 | | 381 | | 48.65 | | | 74.39 |
| M090997-140811A | 540,030 | 385.34 | | 873 | | 387 | | 49.37 | | | 75.59 |
| M090997-140825A | 543,890 | 398.62 | | 897 | | 399 | | 49.42 | | | 75.87 |
| M090997-140906A | 485,655 | 349.44 | | 885 | | 390 | | 49.66 | | | 74.4 |
| M090997-140909A | 483,434 | 350.94 | | 885 | | 396 | | 49.93 | | | 77.04 |
| M090997-140923A | 446,055 | 319.16 | | 879 | | 387 | | 48.24 | | | 76.17 |
| M090997-141004A | 464,184 | 326.39 | | 849 | | 384 | | 48.01 | | | 75.15 |
| M091289-140724A | 436,902 | 318.07 | | 903 | | 390 | | 49.06 | | | 75.31 |
| M091289-140811A | 548,787 | 383.95 | | 852 | | 381 | | 48.24 | | | 74.06 |
| M091289-140825A | 446,864 | 327.73 | | 903 | | 399 | | 50.05 | | | 76.2 |
| M091289-140906A | 529,208 | 381.87 | | 894 | | 387 | | 48.17 | | | 75.44 |
| M091289-140909A | 444,078 | 331.96 | | 927 | | 405 | | 48.84 | | | 78.34 |
| M091289-140923A | 489,180 | 351.84 | | 882 | | 387 | | 47.55 | | | 76.32 |
| M091289-141004A | 499,886 | 354.85 | | 864 | | 387 | | 48.41 | | | 76.42 |
| M101043-140724A | 474,808 | 331.76 | | 858 | | 372 | | 49.14 | | | 74.74 |
| M101043-140811A | 423,615 | 314.89 | | 924 | | 399 | | 50.49 | | | 76.77 |
| M101043-140825A | 469,721 | 335.63 | | 882 | | 381 | | 51.21 | | | 75.87 |
| M101043-140906A | 438,887 | 316 | | 888 | | 384 | | 50.33 | | | 77.35 |
| M101043-140909A | 376,937 | 285.38 | | 939 | | 411 | | 51.42 | | | 78.49 |
| M101043-140923A | 467,518 | 338.58 | | 888 | | 390 | | 49.02 | | | 78.22 |
| M101043-141004A | 436,707 | 319.27 | | 891 | | 399 | | 49.11 | | | 75.67 |
| GeneSet | 3,236,120 | 2,363.70 | | 897 | | 399 | | 47.78 | | | 75.99 |

**Table S3, related to figure 3. The reference gene catalogue of *Macaca mulatta* gut metagenome.**

| **Sample** | **Total Raw Reads (Mb)** | **Total Clean Reads (Mb)** | **Total Clean Bases (Gb)** | **Clean Reads Ratio (%)** | **Total Mapping Ratio (%)** | **Uniquely Mapping Ratio (%)** |
| --- | --- | --- | --- | --- | --- | --- |
| lymph_monkey1_1 | 47.85 | 46.53 | 2.33 | 97.24 | 68.28% | 57.46% |
| lymph_monkey1_3 | 45.41 | 44.09 | 2.2 | 97.1 | 68.70% | 58.24% |
| lymph_monkey1_4 | 44.47 | 43.19 | 2.16 | 97.13 | 68.58% | 57.19% |
| lymph_monkey1_5 | 45.85 | 44.57 | 2.23 | 97.21 | 68.58% | 57.89% |
| lymph_monkey1_6 | 51.14 | 49.66 | 2.48 | 97.11 | 67.12% | 56.21% |
| lymph_monkey1_7 | 38.16 | 37.04 | 1.85 | 97.07 | 67.11% | 55.77% |
| lymph_monkey2_1 | 35.97 | 34.88 | 1.74 | 96.98 | 66.29% | 54.62% |
| lymph_monkey2_3 | 37.86 | 36.67 | 1.83 | 96.85 | 68.67% | 57.89% |
| lymph_monkey2_4 | 34.62 | 33.6 | 1.68 | 97.05 | 68.14% | 56.31% |
| lymph_monkey2_5 | 39.37 | 38.28 | 1.91 | 97.22 | 69.25% | 54.36% |
| lymph_monkey2_6 | 40.95 | 39.77 | 1.99 | 97.12 | 70.53% | 58.94% |
| lymph_monkey2_7 | 40.51 | 39.27 | 1.96 | 96.95 | 64.59% | 51.65% |
| lymph_monkey3_3 | 20.29 | 18.53 | 0.93 | 91.31 | 69.90% | 59.15% |
| lymph_monkey3_4 | 10.54 | 9.61 | 0.48 | 91.23 | 64.84% | 51.30% |
| lymph_monkey3_6 | 22.35 | 20.43 | 1.02 | 91.39 | 69.94% | 58.91% |
| lymph_monkey3_7 | 20.54 | 18.71 | 0.94 | 91.11 | 68.11% | 56.62% |
| lymph_monkey4_3 | 17.85 | 16.17 | 0.81 | 90.63 | 69.41% | 58.59% |
| lymph_monkey4_5 | 17.14 | 15.7 | 0.78 | 91.58 | 67.69% | 52.26% |
| lymph_monkey4_6 | 20.17 | 18.38 | 0.92 | 91.11 | 68.16% | 57.36% |
| lymph_monkey4_7 | 24.02 | 21.94 | 1.1 | 91.33 | 67.99% | 56.41% |
| lymph_monkey5_1 | 17 | 15.58 | 0.78 | 91.64 | 67.03% | 54.35% |
| lymph_monkey5_3 | 19.94 | 18.17 | 0.91 | 91.15 | 69.98% | 59.06% |
| lymph_monkey5_5 | 29.58 | 28.53 | 1.43 | 96.44 | 62.58% | 51.88% |
| lymph_monkey5_6 | 31.76 | 30.65 | 1.53 | 96.49 | 66.06% | 56.16% |
| lymph_monkey5_7 | 25.75 | 24.83 | 1.24 | 96.43 | 62.01% | 52.32% |

**Table S4, related to figure 4. Sequencing data of transcriptome before and after quality control.**

**A**

| **Pathway Name** | **Metabolite count** |
| --- | --- |
| mcc00350 Tyrosine metabolism | 5 |
| mcc01230 Biosynthesis of amino acids | 5 |
| mcc04974 Protein digestion and absorption | 5 |
| mcc00250 Alanine, aspartate and glutamate metabolism | 4 |
| mcc00380 Tryptophan metabolism | 3 |
| mcc02010 ABC transporters | 3 |
| mcc04978 Mineral absorption | 3 |
| mcc00970 Aminoacyl-tRNA biosynthesis | 3 |
| mcc00230 Purine metabolism | 2 |
| mcc00260 Glycine, serine and threonine metabolism | 2 |
| mcc00270 Cysteine and methionine metabolism | 2 |
| mcc00290 Valine, leucine and isoleucine biosynthesis | 2 |
| mcc00760 Nicotinate and nicotinamide metabolism | 2 |
| mcc01210 2-Oxocarboxylic acid metabolism | 2 |
| mcc04024 cAMP signaling pathway | 2 |
| mcc04080 Neuroactive ligand-receptor interaction | 2 |
| mcc04261 Adrenergic signaling in cardiomyocytes | 2 |
| mcc04917 Prolactin signaling pathway | 2 |
| mcc04923 Regulation of lipolysis in adipocytes | 2 |
| mcc04924 Renin secretion | 2 |
| mcc05230 Central carbon metabolism in cancer | 2 |

**B**

| **Pathway Name** | **Metabolite count** |
| --- | --- |
| mcc00380 Tryptophan metabolism | 2 |
| mcc00330 Arginine and proline metabolism | 2 |

**C**

| **Pathway Name** | **Metabolite count** |
| --- | --- |
| mcc00380 Tryptophan metabolism | 4 |
| mcc00350 Tyrosine metabolism | 2 |
| mcc00564 Glycerophospholipid metabolism | 2 |
| mcc04974 Protein digestion and absorption | 2 |
| mcc04723 Retrograde endocannabinoid signaling | 2 |
| mcc00340 Histidine metabolism | 2 |

**Table S5, related to figure 2. Metabolic pathways in muscle, HIP and dmPFC samples, respectively.**

(A) Metabolic pathways in muscle samples.
(B) Metabolic pathways in HIP samples.
(C) Metabolic pathways in dmPFC samples.

| **Genus** | **mean(T1)** | **mean(T2)** | **mean(T3)** | **mean(T4)** | **mean(T5)** | **mean(T6)** | **mean(T7)** |
| --- | --- | --- | --- | --- | --- | --- | --- |
| *Acidovorax* | 3.756E-06 | 1.061E-06 | 0 | 4.654E-07 | 3.519E-07 | 9.517E-06 | 8.842E-06 |
| *Acinetobacter* | 3.353E-03 | 1.679E-05 | 3.143E-05 | 4.842E-05 | 4.262E-04 | 4.658E-03 | 8.904E-03 |
| *Actinobaculum* | 0 | 1.897E-07 | 0 | 0 | 0 | 0 | 0 |
| *Alcaligenes* | 0 | 5.314E-06 | 0 | 3.999E-06 | 0 | 1.806E-07 | 0 |
| *Apibacter* | 1.112E-07 | 0 | 0 | 0 | 0 | 1.922E-07 | 3.556E-08 |
| *Aureimonas* | 0 | 0 | 0 | 2.743E-06 | 0 | 0 | 0 |
| *Bifidobacterium* | 1.905E-04 | 2.207E-04 | 1.661E-04 | 1.833E-04 | 1.896E-04 | 1.463E-04 | 1.611E-04 |
| *Bordetella* | 3.535E-09 | 3.033E-06 | 9.132E-09 | 1.119E-05 | 0 | 1.732E-07 | 1.164E-08 |
| *Cecembia* | 0 | 8.705E-08 | 0 | 8.208E-07 | 6.343E-07 | 0 | 9.404E-07 |
| *Celeribacter* | 0 | 0 | 5.348E-09 | 7.858E-07 | 0 | 7.506E-08 | 0 |
| *Citrobacter* | 3.336E-05 | 1.700E-07 | 2.064E-08 | 1.812E-08 | 2.914E-05 | 1.028E-05 | 1.553E-05 |
| *Comamonas* | 4.119E-04 | 5.165E-07 | 3.153E-07 | 6.075E-07 | 5.287E-05 | 7.202E-04 | 1.384E-03 |
| *Corynebacterium* | 6.306E-05 | 9.826E-04 | 2.639E-05 | 8.824E-06 | 6.361E-05 | 2.120E-06 | 4.655E-06 |
| *Delftia* | 2.039E-06 | 4.618E-09 | 0 | 0 | 1.605E-07 | 4.099E-06 | 6.544E-06 |
| *Desulfitibacter* | 5.645E-07 | 2.197E-07 | 4.133E-08 | 1.022E-07 | 1.242E-07 | 1.159E-07 | 1.586E-07 |
| *Devosia* | 0 | 0 | 0 | 3.928E-07 | 0 | 0 | 0 |
| *Elizabethkingia* | 3.221E-07 | 0 | 0 | 4.661E-08 | 1.786E-08 | 3.605E-07 | 3.465E-07 |
| *Empedobacter* | 7.754E-05 | 1.091E-08 | 7.056E-08 | 5.780E-06 | 1.694E-06 | 1.033E-04 | 4.673E-05 |
| *Exiguobacterium* | 1.272E-07 | 1.957E-08 | 0 | 0 | 0 | 0 | 9.914E-09 |
| *Faecalitalea* | 2.127E-05 | 1.906E-05 | 1.446E-05 | 1.205E-05 | 1.242E-05 | 1.329E-05 | 1.655E-05 |
| *Gordonibacter* | 2.194E-06 | 1.028E-06 | 5.575E-07 | 6.169E-07 | 5.786E-07 | 2.992E-07 | 8.376E-07 |
| *Ignatzschineria* | 1.913E-06 | 5.739E-04 | 1.527E-06 | 1.562E-03 | 0 | 1.042E-08 | 6.687E-07 |
| *Johnsonella* | 4.999E-06 | 7.434E-06 | 5.189E-06 | 6.681E-06 | 5.750E-06 | 3.239E-06 | 3.760E-06 |
| *Klebsiella* | 1.267E-06 | 7.350E-07 | 4.247E-07 | 3.395E-07 | 4.867E-06 | 2.385E-05 | 2.292E-06 |
| *Kluyvera* | 3.392E-06 | 1.192E-07 | 3.567E-08 | 2.988E-08 | 3.476E-05 | 2.412E-05 | 6.871E-06 |
| *Kurthia* | 1.814E-03 | 6.792E-04 | 1.309E-05 | 6.556E-06 | 1.193E-05 | 1.693E-04 | 3.330E-04 |
| *Lactococcus* | 3.687E-05 | 2.515E-06 | 3.894E-06 | 2.341E-06 | 1.064E-04 | 4.763E-04 | 6.861E-04 |
| *Leucobacter* | 6.988E-09 | 1.040E-06 | 2.110E-08 | 1.327E-08 | 0 | 1.400E-09 | 0 |
| *Leuconostoc* | 6.511E-07 | 3.096E-08 | 1.012E-07 | 1.891E-07 | 1.368E-06 | 4.115E-06 | 4.535E-06 |
| *Limnohabitans* | 1.249E-07 | 0 | 0 | 0 | 1.121E-08 | 2.344E-07 | 3.407E-07 |
| *Lysinibacillus* | 2.077E-04 | 6.082E-05 | 3.247E-07 | 1.555E-06 | 6.302E-07 | 1.423E-04 | 2.429E-04 |
| *Macrococcus* | 4.567E-07 | 1.683E-07 | 0 | 1.132E-08 | 0 | 0 | 0 |
| *Marinospirillum* | 0 | 0 | 0 | 1.252E-06 | 0 | 9.592E-08 | 0 |
| *Moraxella* | 2.406E-07 | 0 | 0 | 0 | 1.376E-09 | 1.450E-06 | 7.329E-08 |
| *Morganella* | 6.265E-08 | 0 | 0 | 1.141E-07 | 0 | 0 | 2.106E-07 |
| *Myroides* | 1.298E-04 | 5.264E-06 | 1.407E-06 | 1.523E-03 | 2.911E-06 | 7.912E-04 | 1.999E-04 |
| *Oblitimonas* | 1.989E-07 | 2.641E-06 | 1.575E-07 | 1.776E-04 | 1.661E-08 | 3.638E-07 | 0 |
| *Oligella* | 2.650E-06 | 1.787E-05 | 2.595E-06 | 2.427E-03 | 6.164E-08 | 2.095E-06 | 1.167E-06 |
| *Pectobacterium* | 7.380E-08 | 0 | 0 | 0 | 0 | 6.997E-07 | 1.380E-07 |
| *Pelistega* | 0 | 0 | 0 | 1.198E-06 | 0 | 0 | 0 |
| *Pelobacter* | 0 | 0 | 0 | 2.477E-07 | 0 | 9.677E-08 | 1.227E-07 |
| *Polaromonas* | 1.089E-07 | 0 | 0 | 0 | 9.780E-09 | 2.587E-07 | 2.848E-07 |
| *Proteus* | 2.616E-07 | 1.373E-07 | 0 | 2.170E-06 | 4.421E-08 | 1.521E-06 | 1.289E-06 |
| *Providencia* | 8.233E-05 | 1.072E-05 | 1.783E-07 | 1.614E-04 | 3.871E-06 | 9.531E-05 | 2.867E-04 |
| *Psychrobacter* | 1.643E-06 | 5.028E-08 | 2.202E-08 | 1.396E-05 | 5.020E-09 | 6.189E-07 | 1.551E-06 |
| *Senegalimassilia* | 8.426E-06 | 3.688E-06 | 2.351E-06 | 3.188E-06 | 4.654E-06 | 2.328E-06 | 5.019E-06 |
| *Shigella* | 2.065E-07 | 3.322E-08 | 0 | 1.792E-07 | 2.979E-07 | 4.067E-07 | 2.015E-06 |
| *Sporosarcina* | 3.597E-06 | 1.387E-04 | 0 | 7.809E-07 | 0 | 1.161E-06 | 4.112E-07 |
| *Stenotrophomonas* | 6.264E-07 | 0 | 0 | 0 | 1.241E-07 | 2.328E-07 | 3.677E-06 |
| *Synergistes* | 8.372E-07 | 1.787E-06 | 1.388E-06 | 1.362E-06 | 1.944E-06 | 8.303E-07 | 8.920E-07 |
| *Taylorella* | 0 | 0 | 0 | 5.205E-07 | 0 | 0 | 0 |
| *Variovorax* | 1.281E-07 | 0 | 0 | 0 | 2.485E-08 | 4.315E-07 | 3.180E-07 |
| *Weeksella* | 1.046E-07 | 0 | 0 | 0 | 0 | 3.436E-07 | 5.512E-08 |
| *Weissella* | 8.975E-06 | 0 | 1.119E-08 | 0 | 8.827E-09 | 0 | 2.014E-07 |
| *Wohlfahrtiimonas* | 7.076E-05 | 7.442E-06 | 1.878E-07 | 1.788E-04 | 1.099E-08 | 1.780E-05 | 2.149E-06 |

**Table S6, related to figure 3. The abundance of 55 genera with significantly changed abundance during T1 to T7.**
